# Detecting signals of critical transition in inclusive teaching in biology education: A computational text analysis of *CourseSource*

**DOI:** 10.1101/2023.08.31.555773

**Authors:** Sokona Mangane, Yuhao Zhao, Carrie Diaz Eaton

**Affiliations:** Department of Mathematics, Bates College, Lewiston, ME, USA; Department of Economics and Department of Mathematics, Bates College, Lewiston, ME, USA; Digital and Computational Studies Program, Bates College, Lewiston, ME USA

## Abstract

We examine every Inclusive Teaching section in articles published by the journal *CourseSource* - undergraduate biology teaching materials and related essays - from 2014 to 2022 to understand the evolution of inclusive teaching ideas as expressed through language. We find a rapid shift in attention to inclusive teaching occurs between 2018 and 2019, marked by an increase in section length, increased use of inclusive teaching keywords, and an increase in complexity of ideas in the semantic network. This critical transition occurs before many of the recent events in 2020 which have renewed conversation on equity in education. Some effect is associated with Writing Workshop Faculty Mentoring Networks, which provide participants with authoring resources and support. However, this alone is not enough to explain the observed shift, suggesting that many other structures, conversations, and investments were already providing the fertile ground for educational equity in biology education.

## Introduction

Equity and inclusion has been an ongoing project in STEM postsecondary education. Recent notable funding drivers have included the National Science Foundation’s INCLUDES program, in addition to dedicated programs for underserved states and institutions. However, many critiques, particularly in the post #GeorgeFloyd racial reckoning, have correctly pointed out how little change there has been. In some STEM fields, graduation rates among some minority groups have actually decreased (Newsome 2021 Apr 12). So is STEM education changing, or is it depressingly static?

The study of systems change is a transdisciplinary field, explored from every discipline. Each discipline has reasons for exploring how systems change, from slow change in resilient systems to critical transitions that may “disrupt” systems (Scheffer et al. 2012). As we conceptualize STEM education in a systems change model we can explore how relationships help systems change (e.g. Diaz Eaton et al. 2022), how structures can help systems change (Kezar 2014; Kania, John et al. 2021; Reinholz et al. 2021), and how language changes (Basile and Lopez 2015).

Observing gradual change in a system does not preclude a critical transition. In fact, it may be responsible for neutrally-appearing changes that set the stage for broader shifts to occur (semi-stable states or nearly neutral adaptive landscapes). Therefore, from a theoretical perspective, there is much interest in detecting signals of “tipping points” before they happen (Scheffer et al. 2012). The difference between seeing only gradual change and having hope of a critical transition is also personally important to those who have experienced educational injustice and/or are working towards educational justice in STEM education, like the co-authors.

In this study, we use *CourseSource* as an archive by which to study change over time in inclusive teaching in biology education. We use the term “inclusive teaching” because all *CourseSource* published lesson plans include a section called “Inclusive Teaching” and use capitalization to distinguish the idea from the section of text in the *CourseSource* article. In biology education, the ideas of inclusive teaching are articulated thoroughly in *CBE-Life Science Education’s* Evidence-based Teaching Guide to Inclusive Teaching (Dewsbury and Brame 2019).

We use mixed methods, including computational text analysis and network analysis, to explore how biology education has changed with respect to inclusive teaching over the last 8 years. In this paper, we explore the following questions:

- Are educators devoting more attention to inclusive teaching over time?
- What language are educators using to describe inclusive teaching practices?
- How does the complexity of inclusive teaching language change over time?
- Are there indicators of a critical transition in inclusive teaching?
- What may be driving critical shifts in inclusive teaching language?

## Context

Computational text analysis methods have been applied to a variety of corpora to understand critical shifts and transitions. For example, research on Twitter networks can be used to understand how conversations around climate change and other environmental issues vary and shift in response to political context (Chang et al. 2022). A recent analysis of mathematics texts explored how topics in mathematics are connected to each other or are ordered and have shifted over time (Christianson et al. 2020). In this study, we use the *CourseSource* articles to understand how the community of biology teacher-scholars is shifting with respect to inclusive teaching.

*CourseSource* (coursesource.org) is a journal which publishes teaching activities and lesson plans. It was initially founded to support undergraduate biology education, and its first articles were published in 2014. *CourseSource* is not just a journal for free lesson material, but they are Open Educational Resources (OER) which are hosted on the QUBES Hub platform. This means that anyone can see, download, adapt, use and even reshare their adaptation (Open Education). In addition, all lessons are aligned with Vision and Change or other professional society outcomes and skills, making these lesson plans particularly poised to drive change in the broader STEM education ecosystem (Diaz Eaton et al. 2022).

*CourseSource* has required authors of lesson articles to submit a subsection called “Inclusive Teaching,” but other article types also include the Inclusive Teaching section. This was the primary reason for choosing *CourseSource* as our data source as it represented a focused author reflection on inclusive teaching. We scraped the Inclusive Teaching section from every *CourseSource* article published from 2014 to 2022, creating a dataset of Inclusive Teaching sections (n = 256). Of these, the vast majority (228) were lesson articles. Of the 30 articles excluded from our study due to a lack of Inclusive Teaching sections, only one was a lesson article in 2016. In 2023, *CourseSource* expanded to accept physics articles, but our dataset only represents life science education. The resulting data and R files used in this paper can be found in our GitHub repository.

We, the authors and researchers, approach this from a variety of perspectives, both disciplinary and lived experiences. Collectively we represent identities that are Black, Asian, White, Latinx, International, born in the US, men, women, heterosexual, queer, faculty, students, mathematicians, data scientists, economists, biologists, discipline-based education researchers, and more - too many to completely list. This diversity of perspectives has been complemented by sharing of work outside of our circle with our broader student and educator communities and with *CourseSource* staff and advisory board members to elicit feedback and encourage data sharing. It should be noted that one author is on the advisory board and editorial board for *CourseSource* and occasionally serves as editor or reviewer. While this facilitated access to some data, historical contextualization of trends, and understanding of the writing and publication process, we also relied on collaborating with co-authors outside of biology and seeking external feedback to triangulate key findings.

## A Critical Transition in Inclusive Teaching

We first calculated the length of Inclusive Teaching sections as an indicator of attention to the importance of Inclusive Teaching (Figure 1a). From these, we seem to see a shift in attention in 2019. We observe an increase in skewness and variance, beginning in 2019 and peaking in 2021, typical signals of critical transitions (Scheffer et al. 2012). Grouping 2014 - 2018 (pre-transition in blue) and 2019 - 2022 (transition years in green) together, we see a more clear difference between these stages (Figure 1b). In the pre-transition years, variance ranges from 3369.802 to 8069.429. In the transition years, variance ranges from 13172.153 to 47724.772.

**Figure 1.**
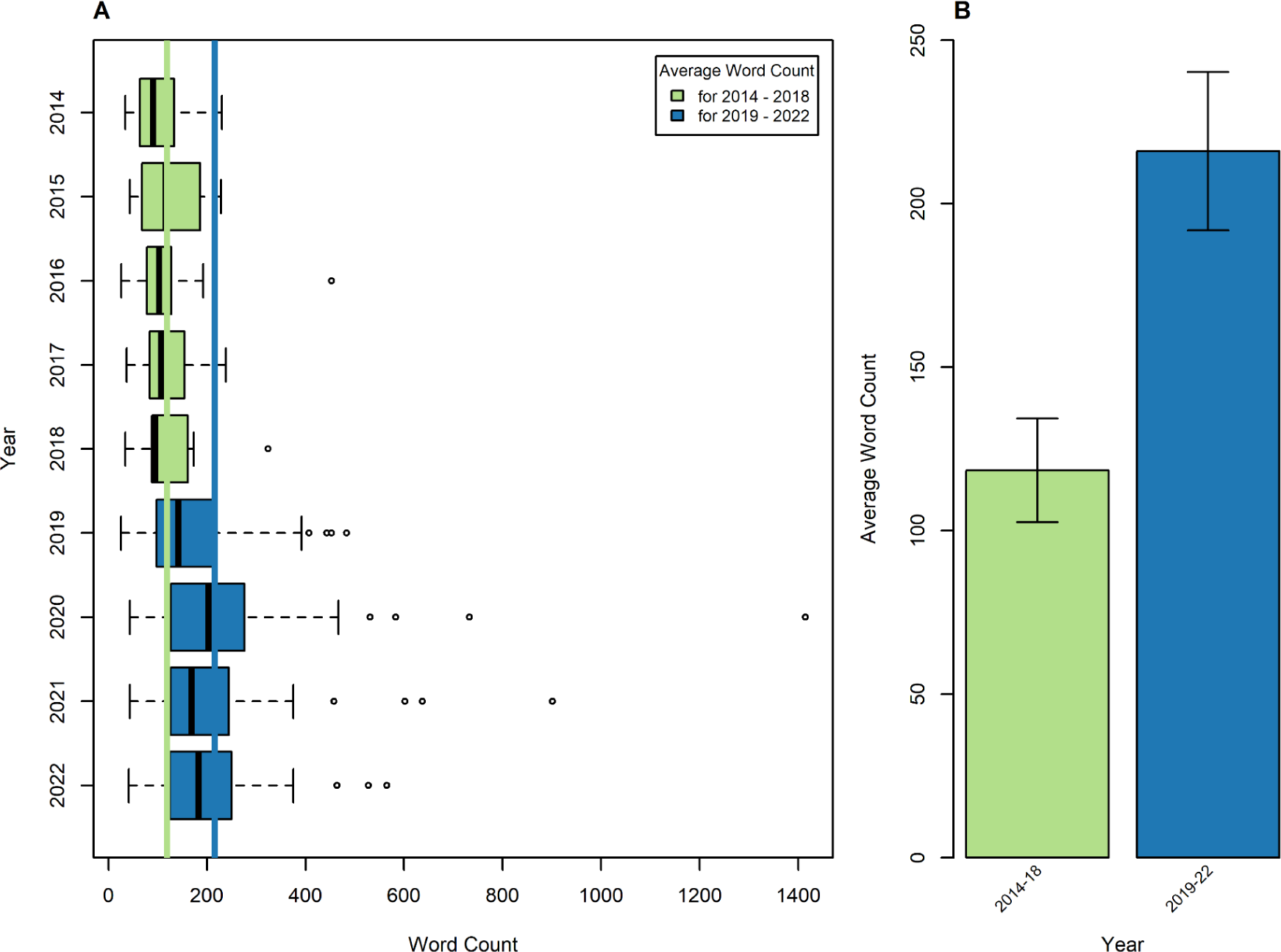
The distribution of inclusive teaching sections each year is plotted in a box and whisker plot in Figure 1a. The green line indicates the average word count between 2014 and 2018 and the blue line is the average word count between 2019 and 2022. Figure 1b displays these means with their confidence intervals indicating a significant difference between groups.

We use the terms transition as opposed to post-transition, because there is not yet enough evidence to indicate whether the transition has concluded or is still in the process of occurring. However, clearly a transition is occurring in attention, and signals of such a transition precede “external shocks” to the system caused by COVID-19 and #GeorgeFloyd - two events often credited with sparking dialogue in inclusive educational practices (e.g. Montgomery 2020; Lemieux et al. 2021). However, from Figure 1a, one may notice that the variance in 2022 begins to subside. This could be due to settling in a new state, and so may be an indication that the transition is nearing a new steady state.

We next examined the actual language in Inclusive Teaching by authors to explore how ideas about inclusive teaching were changing. We cleaned the Inclusive Teaching data by removing stop words (e.g. “and” “or”) and punctuation, and then treating the remaining words as the corpus of interest associated with each article. As a research group, we collectively defined a “JEDI” word as a word associated with social justice, equity, diversity, inclusion, and accessibility. The word could be a subject of the JEDI intervention - e.g. “student”, “environment”, “Hispanic”, indication of instructor design choice - e.g. “designed”, “visual”, “active”, or describe a desired outcome - e.g. “agency”, “diversity”, “conversations”. We were broad in our classification in that we also allowed for all words related to interdisciplinarity connections, multiple representations, social interaction, and active engagement to be classified as JEDI.

Author CDE reviewed each word for a first round classification of JEDI or not. If any word was not immediately clear, a team member would conduct a close reading of the original text. A word was considered JEDI if it (1) was exclusively or almost exclusively used in the context of addressing equity and inclusion and (2) there was no other word in that context that was a much stronger indicator of the JEDI concept. We did not attempt to discern asset-based from deficit-based conceptions of JEDI - for our purposes we wanted all indicators of inclusive teaching ideas. If categorization of a word was still unclear based on the above criteria, we discussed the word as a team and came to a consensus. This process resulted in 534 JEDI words. We generated 2-word, or “duple”, lists from the JEDI words, and repeated this process, resulting in 2431 JEDI duples. The final duples list also includes a few word phrases we identified that may not have a clear JEDI indicator word, but the two-word concept was JEDI, for example “achievement gaps”. A list of the most frequently used JEDI words and duplex can be found in Table 1.

**Table 1:** Top ten most frequent JEDI words and JEDI duples.

| Word | Count | Duple | Count |
| --- | --- | --- | --- |
| students | 1389 | active learning | 38 |
| inclusive | 123 | inclusive teaching | 36 |
| diversity | 115 | learning environment | 24 |
| diverse | 100 | group members | 21 |
| opportunity | 94 | help students | 18 |
| individual | 72 | encourages students | 15 |
| engage | 66 | inclusive learning | 15 |
| environment | 64 | community college | 13 |
| background | 57 | encourage students | 11 |
| participate | 57 | self-efficacy | 11 |

Consistent across all years is the use of the words “students” and “inclusive teaching.” However, when comparing frequency of JEDI word use across years, we find similar signals of transition from 2018 to 2019 (Figure 2a). Figure 2a is log scaled due to the dominance of the word “students” and thus dampens the appearance of difference. Even so, we can still see that in the pre-transition years, there is less variety of language use and inconsistency in which language is most used. In the transition years, there are a lot of different JEDI words being used and more consistently across years.

**Figure 2.**
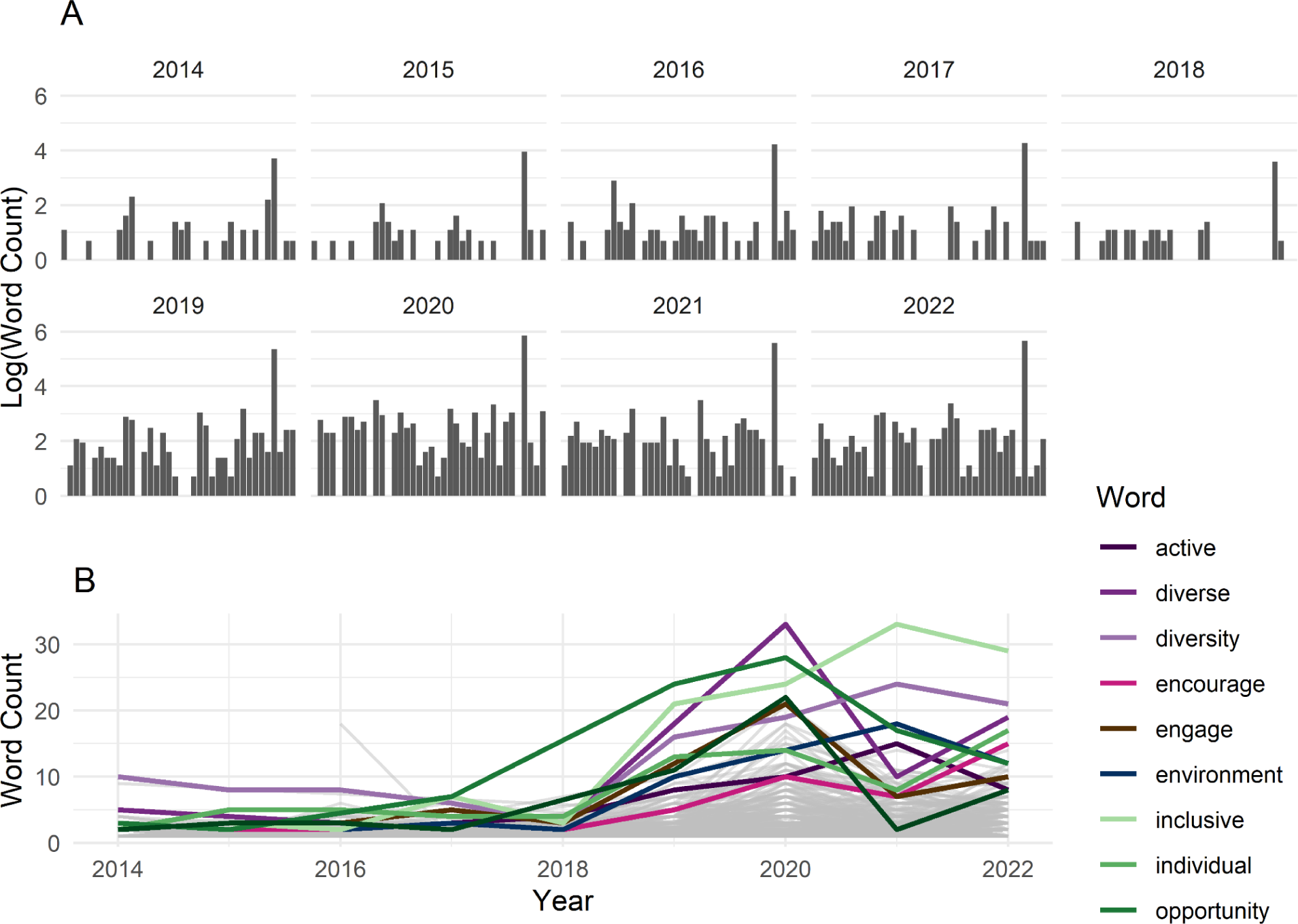
The log of the word frequency over time is plotted in Figure 2a. This is the word frequency of the top 40 words across all years. Figure 2b displays the word frequency over time of the top 9 words after 2018. Figure 2b excludes the most used word, “students”.

We can also examine the trends of individual words over time (Figure 2b), and notice some differences in the language use that is occurring. After 2018, we see an increase in and maintenance of the word “inclusive” - likely no surprise as the name of the section is Inclusive Teaching, and it is one of the most frequently used JEDI duples as well (Table 1). However, note that “inclusive learning” also appears frequently. Unlike “inclusive teaching” which is both associated with the section name and the aforementioned

## Inclusive teaching language complexity

We wanted to understand whether the inclusive teaching shift we detected was performative - just an increased inclusive word use, or if there was a signal of more complex understanding of inclusive language through complex language development. Across all articles, approximately 90% of the variation in JEDI word count in each article can be explained by the overall length of its Inclusive Teaching section. When we examined JEDI word “density” of an article - the number of JEDI words divided by the total number of words in the Inclusive Teaching section - we continued to find a consistent shift between 2018 and 2019. However, the years with the highest average density across articles were the first two years of journal publication, 2014 and 2015, and in these years JEDI density was 0.21 and 0.22, respectively.

Another way to think about complex JEDI ideas is to look at JEDI phrases. For this analysis we treated our corpus as a semantic language network. Semantic networks are language networks built from a set of words that convey particular ideas (Solé et al. 2010). We used the JEDI duple list constructed from the JEDI list to construct a directed word network for each year. Each word in the JEDI duple is a node, with the arrow indicating the order of the word combination. From Figure 3, we can see that the network gains considerable complexity over time. The increased complexity is visible through the acquisition of more nodes over time. Also, ideas which were once communicated through separate networks on the periphery are becoming connected to the main idea, or the largest subgraph.

**Figure 3.**
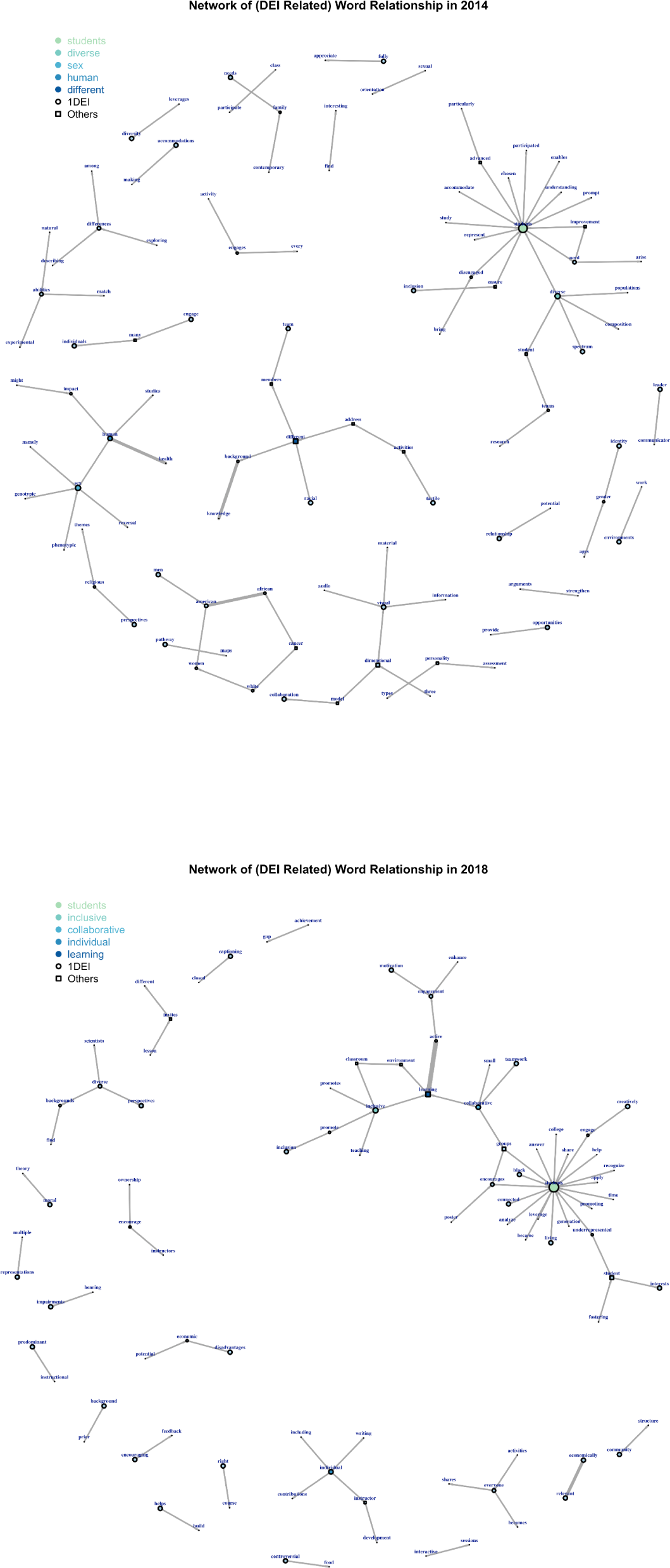
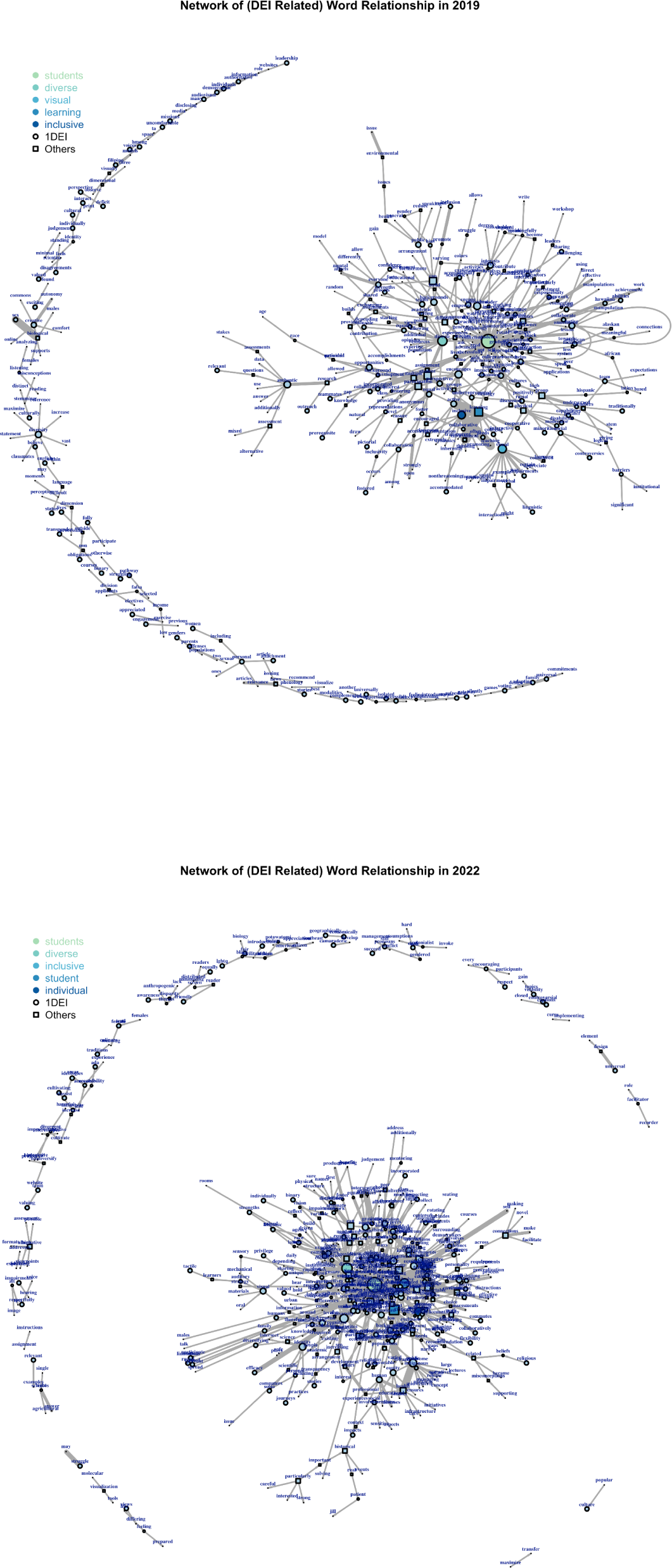
The semantic network in 2014, 2018, 2019, 2022. All four semantic networks plotted using edge-lists(two word pairs)involving the JEDI words we are focusing on. Critical transition is reflected in network change from 2018-2019.

In semantic network studies, complexity of language is measured through typical measures of network complexity (Christianson et al. 2020). These include network size (number of nodes in a network), edge density (ratio of edges present to number of edges possible), diameter (the maximum of all possible shortest path lengths between two nodes), clustering coefficient (relative number of triangles in the network graph), number of disjoint components, and size of the largest component. In Table 2, see size, diameter, number of components, and maximum component size experience a jump and that increase points to a critical transition. Notice that the component is defined as the subgroup in networks in which all actors are connected, directly or indirectly.

**Figure 4.**
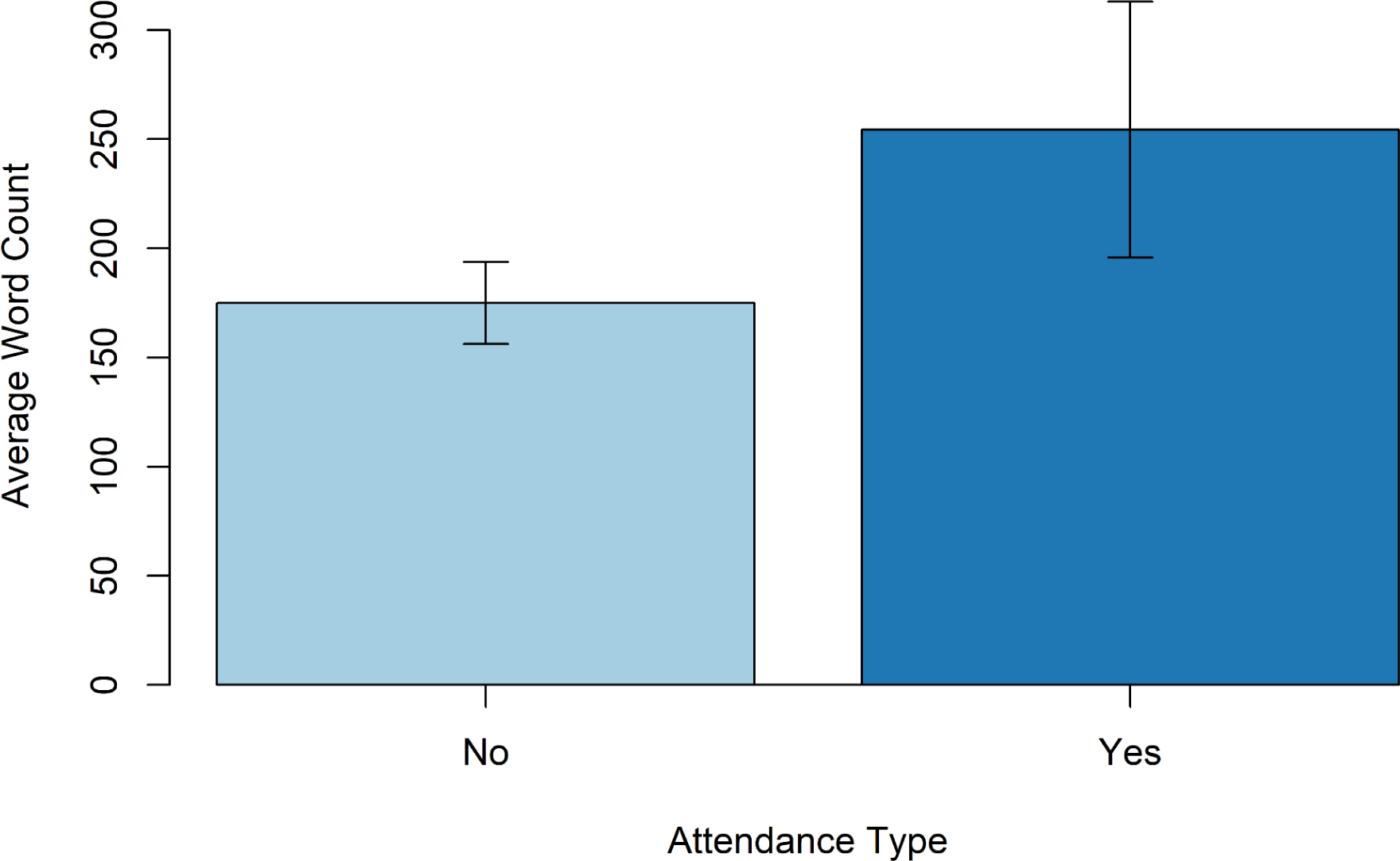
The average word count of the Inclusive Teaching section, with confidence intervals, for those with at least one author who attended a Writing Workshop FMN (Yes - dark blue) and those that did not (No - light blue).

**Table 2:** Language network measure summary. The green shades the pre-transition 2014 - 2018 years and the blue shades the 2019 - 2022 transition years.

| Year | Size | Edge Density | Diameter | Cluster Coeff | #Components | Max comp size |
| --- | --- | --- | --- | --- | --- | --- |
| 2014 | 108 | 0.007355486 | 5 | 0.018072289 | 25 | 25 |
| 2015 | 160 | 0.004795597 | 8 | 0.006012024 | 43 | 50 |
| 2016 | 164 | 0.004788269 | 5 | 0 | 38 | 46 |
| 2017 | 160 | 0.004992138 | 6 | 0 | 38 | 69 |
| 2018 | 92 | 0.008719541 | 4 | 0.013157895 | 21 | 40 |
| 2019 | 447 | 0.002207040 | 14 | 0.013135736 | 59 | 300 |
| 2020 | 663 | 0.001722464 | 17 | 0.014102564 | 50 | 537 |
| 2021 | 496 | 0.002122027 | 14 | 0.009406355 | 49 | 381 |
| 2022 | 585 | 0.001765016 | 12 | 0.011164274 | 66 | 423 |

The network in pre-transition years has a higher connectivity relative to transition years. The connectivity is an indicator of critical transition as networks with higher connectivity generally change more abruptly (Scheffer et al. 2012). Connectivity likely experienced a large drop because the semantic network experienced a large influx of new language nodes in 2019. In 2019 and on, the connectivity is much smaller, which may indicate that change, moving forward, is more likely to occur relatively gradually (Scheffer et al. 2012).

## Why is Inclusive Teaching Language Shifting?

We were curious about why inclusive language was experiencing a shift, particularly since the shift is detected before 2020 when such teaching shifts are attributed often to the “external shocks” of COVID-19 and #GeorgeFloyd. *CourseSource* has run writing workshops, and in a data set shared with us, it was typically about a year from workshop date to final publication. That means that the influence of events in 2020 would most likely be detectable beginning in 2021. What we are seeing here is an earlier influence, circa 2018 submission and likely earlier implementation in the classroom.

Within STEM Education, there were some notable events in 2017 and 2018 related to funding investments in science education. In 2017, the Howard Hughes Medical Institute (HHMI) launched its Inclusive Excellence program, asking 24 colleges and universities to transform its practices and improve graduation rates among students that institutions had been failing (HHMI 2023). In 2018, the National Science Foundation (NSF) released the call for their INCLUDES program (Inclusion across the Nation of Communities of Learners of Underrepresented Discoverers in Engineering and Science), which focused on broadening participation through research and support to scale (2018 Jan 5). Finally, CourseSource received a grant in 2017 to support writing workshops, run as online Faculty Mentoring Networks in collaboration with QUBES (www.qubeshub.org). These Writing Workshop FMNs launched in 2018, which also began their data collection on workshop attendees, with the first Writing Workshop FMN participant publishing in 2019.

*CourseSource* allowed us to access a list of articles in which at least one author had attended their Writing Workshop FMN. Because we already knew that there was a significant shift in 2019, we limited our comparison to articles from the same time frame, 2019 to 2022. We found a significant and meaningful difference (p-value = 0.012) in the average length of the Inclusive Teaching section between these two time intervals (see Figure 3). The average JEDI word count illustrates a similar pattern (p-value = 0.016).

Authors are devoting more attention to the Inclusive Teaching section and the development of writing workshops which support authors in writing for *CourseSource* are associated with this increasing attention. When sharing preliminary results with *CourseSource*, we learned that during workshops, participants receive example articles, are pointed to example Inclusive Teaching sections, and are given some examples of topics and resource articles to consider when writing their Inclusive Teaching section. This is not exclusive to the Inclusive Teaching, but this guidance is applied to all sections. However, the difference between the workshop participants’ articles and the non-workshop participants’ articles is much smaller than the difference between the pre-transition and transition years. This indicates that there are likely other factors influencing the broader shift we are seeing.

## Implications

The shift in the number and variety of words to describe inclusive teaching may be expected in a critical transition (Scheffer et al. 2012), but there are practical implications. As we try out new words, their meanings and intentions may not yet be universally known and established. In addition, it is more difficult to locate inclusive teaching material or ideas if there is not a consistent set of words used to describe it or links made through metadata (Piedra et al. 2017). Having a dedicated author-written Inclusive Teaching section becomes extremely useful metadata. This section gives both readers and researchers the ability to make the connection between inclusive teaching language and the intent in an instructor talk context. In addition, having an entire Inclusive Teaching section may be more effective than having a controlled vocabulary alone, particularly while the language of the field is shifting.

Some of the increase in JEDI word count use may be related to author participation in writing workshops. It may be worth following up with co-authors to understand how writing workshop FMNs may have influenced their articulation of inclusive teaching. However, while this may account for some of the variance within 2019 to 2022, the structured professional development experience alone is not enough to explain the shift observed after 2018.

It is likely that years of work have set the conditions and built the capacity to make shifts in inclusive teaching. As noted above, HHMI and NSF are among those that have made significant monetary investments in inclusive postsecondary STEM education in the US. In addition to research and professional development funded by these initiatives, there have been a multitude of other conversations which were likely crucial to triggering the shift we observed. It is worth noting that continuing these investments and conversations will be critical (e.g. She-shonda Porter et al. 2023). The decrease in variance from 2021 to 2022 seen in Figure 1a may be indicative of a settling into a new steady state and a shift into more gradual change.

As scholars of inclusive teaching, sometimes we feel “And still we see no change” (Basile and Lopez 2015), and indeed the statistics from 2014 - 2018 look depressingly static. However, this study provides some indication that change is possible, and that already the “fossil record” of our curricular archives are already showing signals of critical transition and rapid change. Detecting these signals gives us hope, and should reinvigorate efforts to continue working towards educational justice for STEM.

## Acknowledgments

This material is based upon work supported by the William and Flora Hewlett Foundation’s support of the RIOS Institute (#2020-1363). We thank *CourseSource* staff and advisory board for their feedback, and their willingness to share data with our research team. We also thank the entire leadership team of the RIOS Institute for feedback on this project, particularly Naz Fruits for helping with the initial JEDI word classification process.

## Author Biographical

Sokona Mangane is an alumna of Bates College in Lewiston, Maine, who majored in Mathematics and minored in Digital and Computational Studies. She is an incoming Data Analyst at Ayers Saint Gross, an architectural firm based in Maryland, which specializes in master plans and building designs for colleges, universities, and cultural facilities.

Yuhao Zhao is a rising senior of Bates College in Lewiston, Maine who majored in Mathematics and Economics and minored in Digital and Computational Studies. He is a network enthusiast and interested in research of intersection of applied math and social sciences.

Carrie Diaz Eaton is an associate professor in the interdisciplinary Digital and Computational Studies Program at the Bates College in Lewiston, Maine and a Principle Investigator and Co-Director of the Institute for a Racially-just, Open, and Inclusive STEM Education (RIOS Institute).

## Data Repository

https://github.com/INQUIRE-Lab/CourseSource

